# P2X1 selective antagonists block HIV-1 infection through inhibition of envelope conformation-dependent fusion

**DOI:** 10.1101/783464

**Authors:** Alexandra Y. Soare, Hagerah S. Malik, Natasha D. Durham, Tracey L. Freeman, Raymond Alvarez, Foramben Patel, Namita Satija, Chitra Upadhyay, Catarina E. Hioe, Benjamin K. Chen, Talia H. Swartz

## Abstract

Purinergic receptors detect extracellular ATP and promote inflammatory processes. Emerging literature has demonstrated that inhibition of these proinflammatory receptors can block HIV-1 productive infection. The specificity of receptor type and mechanism of interaction has not yet been determined. Here we characterize the inhibitory activity of P2X1 receptor antagonists, NF279 and NF449 in cell lines, primary cells, and in a variety of envelope clades. NF279 and NF449 blocked productive infection at the level of viral membrane fusion with a range of inhibitory activities against different HIV-1 envelopes. A mutant virus carrying a truncation deletion of the C-terminal tail of HIV-1 envelope (Env) glycoprotein 41 (gp41) showed reduced sensitivity to P2X1 antagonists, indicating that the sensitivity of inhibition by these molecules is modulated by Env conformation. By contrast, a P2X7 antagonist, A438079, had limited effect on productive infection and fusion. Inhibition with NF449 interfered with the ability of the V1V2 targeted broadly neutralizing antibody PG9 to block productive infection, suggesting that these drugs may antagonize HIV-1 Env at gp120 V1V2 to block viral membrane fusion. Our observations indicate that P2X1 antagonism can inhibit HIV-1 replication at the level of viral membrane fusion through interaction with Env. Future studies will probe the nature of these compounds in inhibiting HIV-1 fusion and in development of a different class of small molecules to block HIV-1 entry.

**IMPORTANCE:** While effective treatment can lower the severe morbidity and mortality associated with HIV-1 infection, patients infected with HIV-1 suffer from significantly higher rates of non-communicable comorbidities associated with chronic inflammation. Emerging literature suggests a key role for P2X1 receptors in mediating this chronic inflammation but the mechanism is still unknown. Here, we demonstrate that HIV-1 infection is reduced by P2X1 receptor antagonism. This inhibition is mediated by interference with HIV-1 Env and can impact a variety of viral clades. These observations highlight the importance of P2X1 antagonists as potential novel therapeutics that could serve to block a variety of different viral clades with additional benefits for their anti-inflammatory properties.

## Introduction

HIV-1 infection represents a major public health concern worldwide with 37 million individuals living with the disease and nearly two million new infections occurring each year (1). Due to the advent and improvement of antiretroviral therapy (ART), people with HIV-1 (PWH) now benefit from sustained suppression of viremia and greatly improved life expectancy. Despite the success of modern ART, longitudinal studies suggest that HIV-1 infection is characterized by chronic inflammation, despite viral suppression (2-9). Despite these challenges, emerging literature has supported a role for the immune signaling receptors known as purinergic receptors for HIV-1 associated inflammation (10-15). Purinergic receptors recognize extracellular nucleotides that are released from inflamed or dying cells and are upstream of a wide variety of signaling pathways that mediate pro-inflammatory responses, including pyroptosis (16, 17). There are several purinergic receptor subtypes that differ in ligand selectivity and structure but share a similar purpose of mediating important physiological cell responses due to environmental cues. These receptors are ubiquitously expressed on mammalian cells and are found on many immune cell subsets important in HIV-1 infection, including lymphocytes and myeloid cells (18-21). Our laboratory and others have explored the role of purinergic receptors in HIV-1 infection and have reported that non-selective purinergic receptor antagonists can block HIV-1 infection and can reduce inflammatory cytokine production associated with infection (22-24).

Of note, the P2X receptor subtype in particular has been explored in the context of HIV-1 pathogenesis (22-24). P2X receptors are nonselective cation channels found on a wide variety of immune cells and are divided into seven subfamilies. Within those subfamilies, the P2X1 and P2X7 receptors are predominantly expressed on T helper cell (T_h_ cells), the primary target of the HIV-1 virus (42, 43). Recently, our laboratory demonstrated that P2X1 and P2X7 antagonists can inhibit HIV-1 infection and inflammatory cytokine production in human tonsil cells, implicating their potential as novel HIV-1 therapeutics. However, the manner and site by which these antagonists are able to block HIV-1 infection has not been fully determined. Inhibition of these receptors through various antagonists has been demonstrated to impact on the HIV-1 life cycle at the stage of viral membrane fusion (22, 23, 25-27). Giroud et al. demonstrated that P2X1 antagonists blocked HIV-1 fusion by blocking virus interactions with co-receptors C-C chemokine-receptor 5 (CCR5) and CXC chemokine-receptor 4 (CXCR4) (22, 27). The authors observed that NF279 could bind and block receptor signaling and arrest HIV-1 fusion downstream of CD4 binding prior to engagement of coreceptor. Our data implicate early HIV-1 fusion events and we sought here to determine the mechanism of fusion inhibition by probing the role of HIV-1 envelope (Env) interaction with the P2X1 antagonists.

Here, we examined the activities of P2X1 antagonists on HIV-1 productive infection in order to determine the impact on HIV-1 fusion, the impact on neutralization of HIV-1 expressing various Env sequences, and the site of localization on HIV-1 Env where these drugs inhibit. We demonstrate that two drugs selective for P2X1, NF279 and NF449, can neutralize a diverse panel of HIV-1 viruses with differing IC_50_ values and that treatment with NF449 can alter Env accessibility and impact the access of broadly neutralizing antibodies, specifically targeting the V1V2 region of HIV-1 Env. These data indicate that P2X1 antagonists can interfere with HIV-1 Env directly, potentially altering the binding of HIV-1 and inhibiting interaction with coreceptor (CCR5/CXCR4) that mediates HIV-1 fusion. This study provides the compelling suggestion that P2X1 antagonists could be used as an adjuvant strategy to alter HIV-1 envelope accessibility in the interest of vaccine and therapeutic development.

## Results

### NF279 and NF449 block HIV-1 productive infection in a lymphocyte cell line in a dose-dependent manner

A panel of antagonists was tested for dose-dependent inhibition of infection of HIV-1 NL-CI (NL4-3 Cherry internal ribosome entry site), an X4-tropic virus with the fluorescent mCherry protein expressed in place of Nef (28). Azidothymidine (AZT), the nucleoside reverse transcriptase inhibitor, and pyridoxalphosphate-6-azophenyl-2′,4′-disulfonic acid tetrasodium salt (PPADS), a non-selective P2 antagonist that we observed previously to inhibit HIV-1 infection in MT4 cells (23), were compared to a variety of P2X1 and P2X7 selective antagonists in the MT4 cell line (Figure 1, Table 1). IC_50_ inhibitory values are indicated where applicable. The most compounds in this model were P2X1 antagonists NF279 and NF449, which inhibited HIV-1 at IC_50_ values of 0.88 μM and 0.23 μM, respectively, as compared with AZT, which had an IC_50_ value of 0.01 μM. Both NF279 and NF449 were more potent than PPADS (IC_50_ = 4.7 μM). A438079 is a P2X7 inhibitor that did not demonstrate inhibitory capability ≥ 50%. Other P2X7 selective drugs including A804598, A839977, AZ11645373, A740003, AZ10606120, GW791343, JNJ479655567, and Ro 0437626 did not inhibit NL-CI infection in MT4 cells. For almost all compounds, minimal toxicity was observed with the exception of 10-20% cell toxicity resulting from 100 μM NF279 treatment and >50% toxicity resulting from 100 μM AZ10606120 or Ro 0437626 treatment.

**Table 1.**
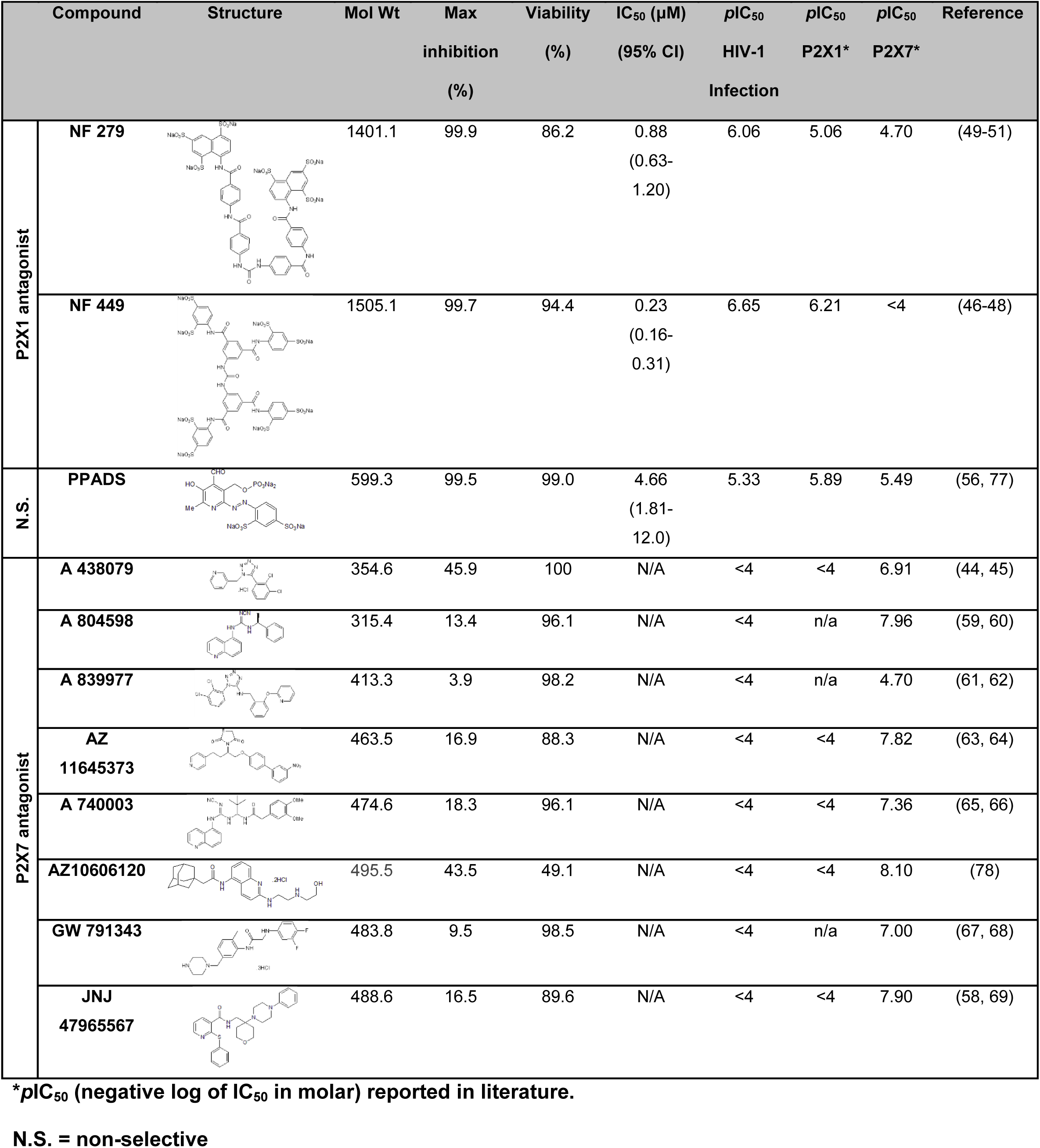
Inhibition of HIV-1 productive infection by P2X-selective antagonists.

**Figure 1.**
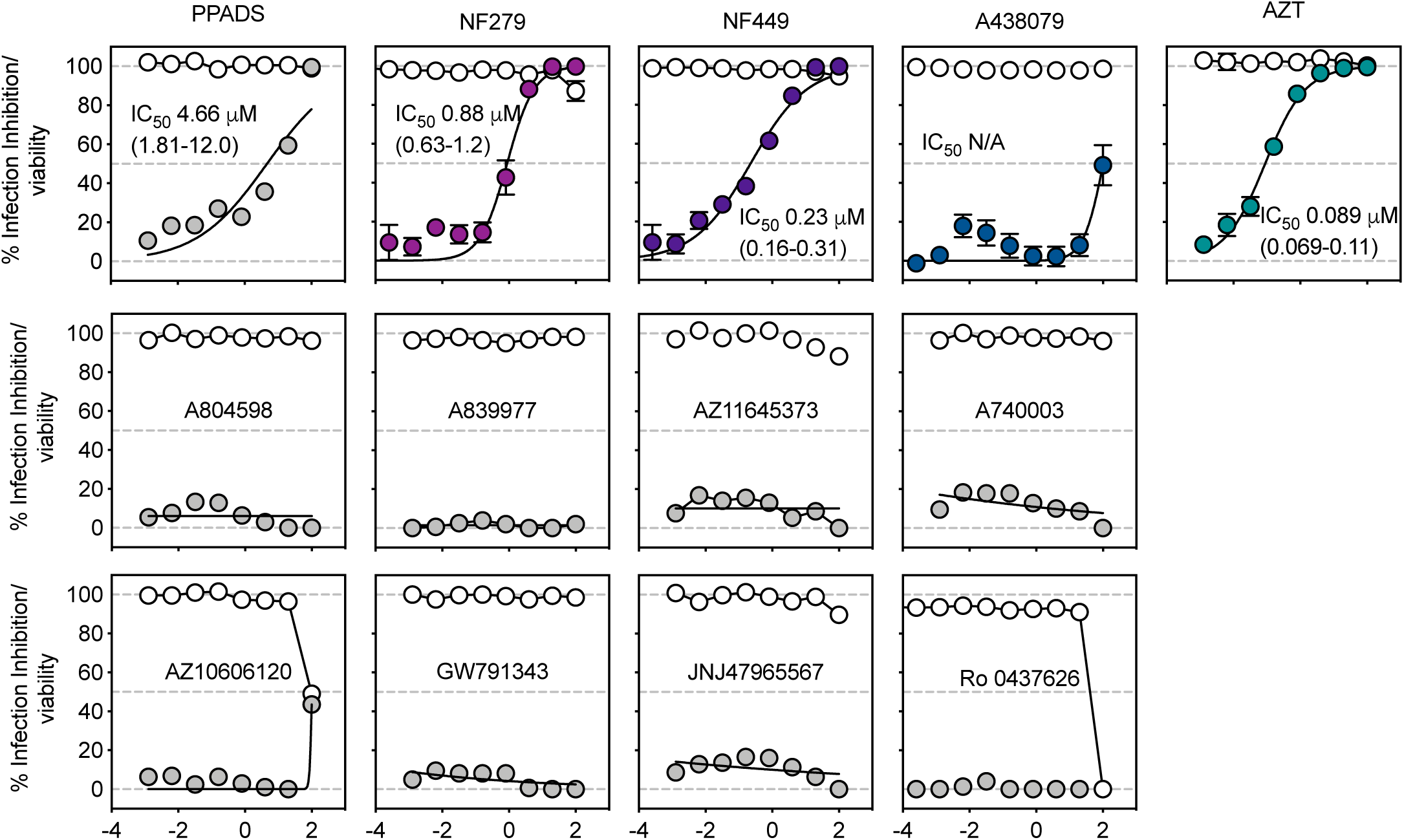
P2X antagonists NF279 and NF449 block HIV-1 productive infection in a dose-dependent manner in MT4 cells. A panel of P2X antagonists was tested for inhibition of productive infection in MT4 cells. Inhibitors were added at indicated concentrations to MT4 cells in the presence of HIV-1 NL-CI (X4-tropic). Nonlinear regression curve fits are shown. Infections were conducted in the presence of 5-fold dilutions of inhibitors from 100 μM. Samples were incubated for 48 hrs, fixed and analyzed by flow cytometry. Inhibition is calculated based on the infection in the presence of drug as a function of the infection without inhibitor. Viability is plotted in the unfilled circles and percent inhibition is plotted in the black/filled circles. IC_50_ values (mean and range) are indicated. Results are the means ± SEMs of at least three independent experiments.

### Kinetics of NF279 and NF449 inhibition confirm their activity on HIV-1 Env-mediated fusion

Our previous studies implicated NF279 and other P2X antagonists as HIV-1 fusion inhibitors in cell-to-cell and cell-free infection (23, 25). Here, we assessed whether the above antagonists were similarly active as HIV-1 fusion inhibitors. We employed a HIV-1 Gag-iCre cell-based viral entry assay, in which Cre-recombinase is packaged into the HIV-1 virion and target indicator cells undergo a Cre-activated red-to-green (RG) switch upon HIV-1 fusion (25, 26). We observed that both NF279 and NF449 inhibited 100% HIV-1 fusion at 100 μM, while A438079 partially inhibited fusion at 100 μM (Figure 2A). Dose response curves indicated a dose-dependent inhibition of HIV-1 fusion by NF279 and NF449, while partial inhibition by A438079 was below 25% for all concentrations tested (Figure 2B).

**Figure 2.**
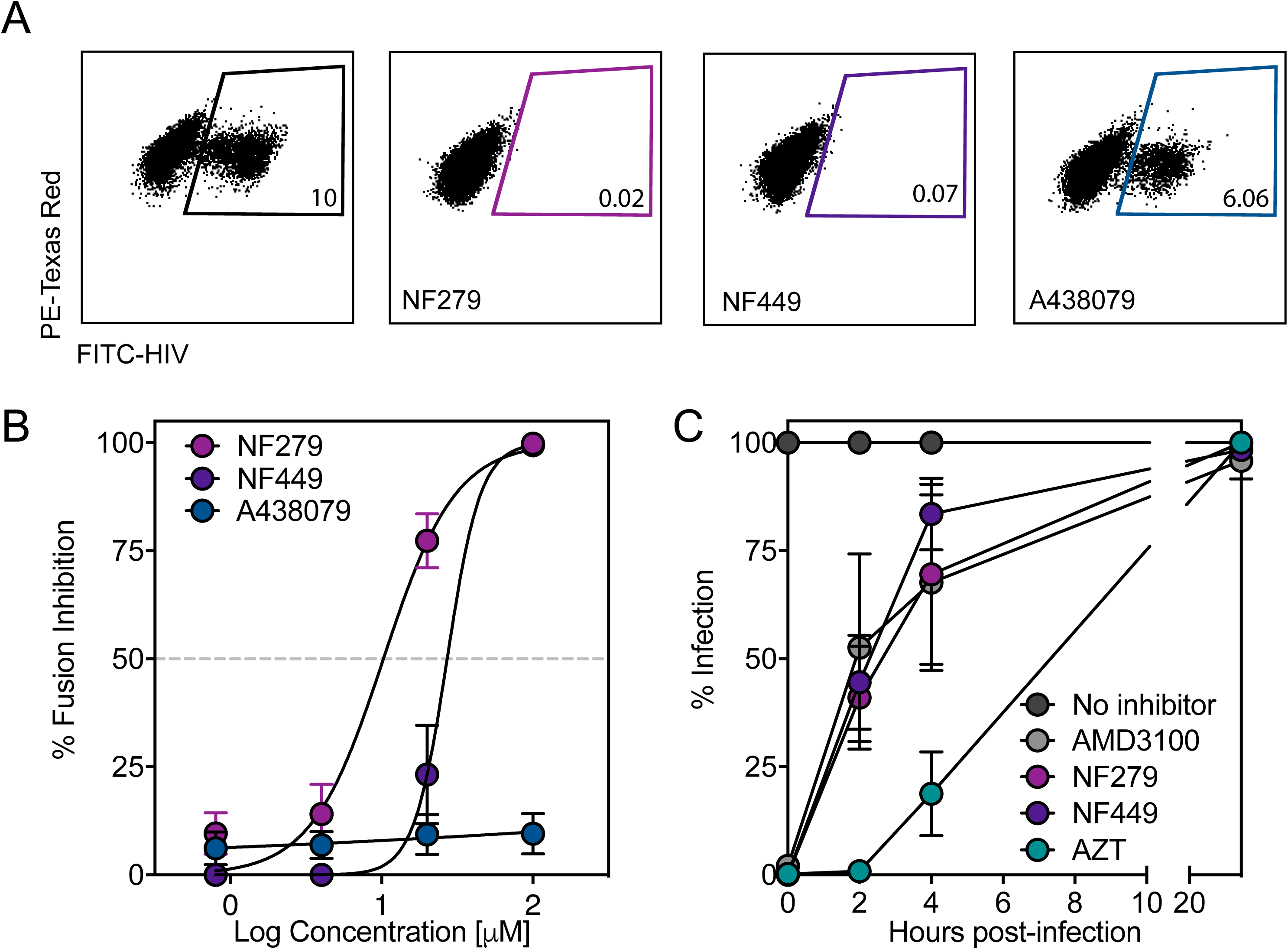
NF279 and NF449 function contemporaneous with HIV-1 fusion inhibitors. (A) Donor Jurkat cells were nucleofected with HIV-1 Gag-iCre and co-cultured with Jurkat RG cells for 48 hrs. RG switch, as a marker of HIV-1 viral membrane fusion, was measured by flow cytometry in the presence or absence of the indicated inhibitors (100 μM) and quantified based on GFP fluorescence. (B) Dose response curves are plotted from the assay described in A for the three compounds: NF279, NF449, A438079. (C) Time-of-addition experiment was performed by measuring productive infection of HIV-1 NL-CI in MT4 cells with inhibitors added at indicated times post-infection. Infection is quantified by flow cytometry and plotted as a function of percentage of total infection without inhibitors at the indicated time points. Results are the means ± SEMs of at least three independent experiments.

We next performed time-of-addition experiments (Figure 2C) in which these drugs were added at different hours post-infection (HPI) to verify whether they acted on the virus fusion step or later stages in the viral life cycle (29, 30). Virus attachment and virus-cell fusion were expected between 0-2 HPI, reverse transcription was expected at 3-4 HPI, integration was expected near 6 HPI, transcription, RNA export, translation and assembly were expected between 10-16 HPI, and budding and maturation were expected after 16 HPI (29, 30). We hypothesized that the P2X1 antagonists would inhibit early viral life cycle events contemporaneous with AMD3100, a CXCR4 antagonist that inhibits viral membrane fusion at 2 HPI. In contrast, AZT is known to act on reverse transcription and was expected to inhibit HIV-1 productive infection when added prior to 4 HPI. MT4 cells were infected with HIV-1 NL-CI and drugs were added at 0, 2, 4, and 24 HPI. We observed that several P2X1 antagonists did not inhibit or weakly inhibited including A438079 and A804598 (not shown), while AMD3100 inhibited at the same time-point as NF279 and NF449, confirming that these drugs inhibit viral replication at the stage of viral membrane fusion. Addition of NF279, NF449, or AMD3100 after 4 HPI did not inhibit infection, as HIV-1 fusion would have already occurred. As expected, addition of AZT prior to 4 HPI resulted in near complete inhibition of infection, while addition of AZT after 4 HPI did not result in inhibition of infection, as reverse transcription had already occurred. We concluded that NF279 and NF449 block HIV-1 infection at same stage of the HIV-1 virus life cycle as the fusion inhibitor, AMD3100.

### Contribution of gp41 to HIV-1 inhibition by NF279 and NF449

It has been hypothesized that P2X1 or P2X7 antagonism through receptor interaction might represent the mechanism of fusion inhibition. Our laboratory and others have demonstrated a dearth of receptor expression on the surface of infected cells as well as a body of data suggesting interaction and activation of coreceptor (22, 27, 31). We therefore probed the interaction of drug directly on HIV-1 envelope.

HIV-1 viral membrane fusion is mediated by HIV-1 envelope glycoproteins gp120 and gp41. Binding of gp120 to CD4 and coreceptor activates a fusogenic function of gp41 by exposing the N-terminal fusion peptide which allows for cell membrane insertion and generation of a pre-hairpin intermediate structure that bridges viral and target cell membranes ((32-37). Given our previous results which suggests that NF279 and NF449 block HIV-1 fusion, we tested whether they interacted with HIV-1 Env gp41. First, we examined whether the role of gp41 pre-hairpin structure was required for fusion inhibition by NF279, NF449, and A438079. To test this, we used enfuvirtide (also known as T-20), a 36-residue alpha peptide fusion inhibitor that binds to the pre-hairpin structure of gp41, thereby inhibiting viral membrane fusion (32-34, 38, 39). We assessed the role of pre-treatment with T-20 on inhibition of productive infection by NF279, NF449, and A438079. We reasoned that T-20-abrogation of the effect of NF279, NF449, or A438079 inhibition would indicate a role for the gp41 hairpin structure in the nature of this interaction. Figure 3A shows the effect of each drug on inhibition of HIV-1 productive infection in the presence (colored bars) or absence (50% opacity colored bars) of T-20 is shown. T-20 was used at a concentration (0.1 μg/ml) that on its own caused 50% inhibition without added drug. T-20 treatment resulted in some inhibition at NF279 0.5 μM and 1 μM but not at 5 μM and this did not mean significance. No effect was noted for NF449 or A438089 or for AZT as would be expected. This suggested that NF449 and likely NF279 antagonism is not dependent on the pre-hairpin structure of gp41.

**Figure 3.**
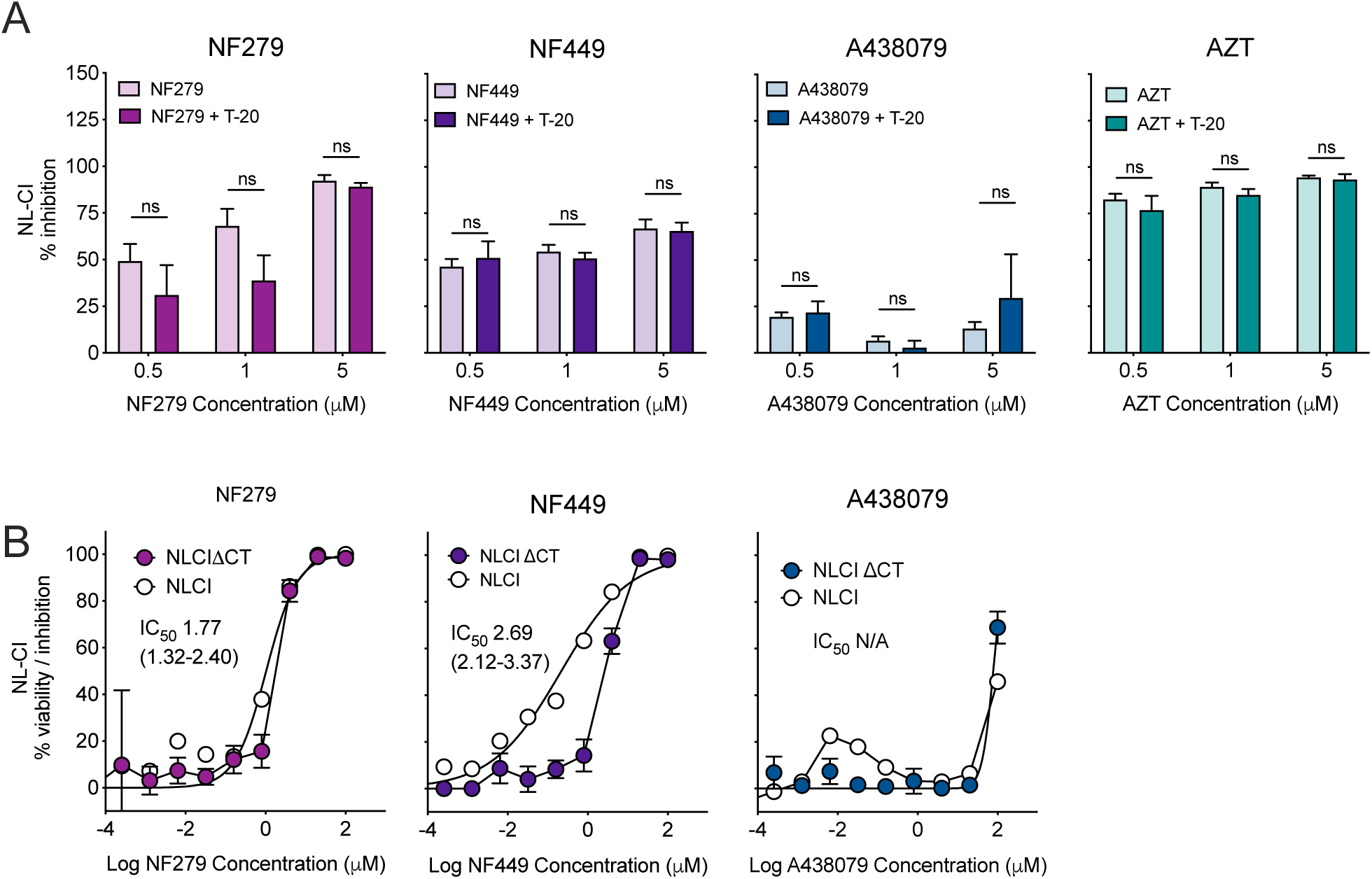
Inhibition of HIV-1 membrane fusion by P2X-seclective antagonists occurs through interactions with HIV-1 Env and not by antagonism of gp41. (A) The effect of T-20 (gp41 hairpin structure inhibitor) on inhibition of HIV-1 NL-CI was tested by pre-incubation of MT4 cells with T-20 and the addition of 5-fold dilutions from 5 μM of the indicated inhibitions of NF279, NF449, A48079 and AZT. Mean values ± standard errors of the means are presented from three donors. ns = not significant. (B) NF279, NF449, and A438079 were added at indicated concentrations to MT4 cells in the presence of HIV-1 NL-CI or NL-CIΔCT. Nonlinear regression fits for cell-free infection in NL-CI (top) and NL-CIΔCT (bottom) are shown. Viability is plotted in the unfilled circles and percent inhibition is plotted in the black circles. Infections were conducted in the presence of 5-fold dilutions of inhibitors from 100 μM. Samples were incubated for 48 hrs, fixed and analyzed by flow cytometry. Results are the means ± SEMs of at least three independent experiments. IC_50_ values are indicated.

Subsequently, we tested the P2X1 antagonists for inhibition of infection in MT4 cells by a C-terminal truncation of HIV-1 Env gp41 (NL-CIΔCT). This mutant enhances Env fusogenicity and has been shown to regulate exposure of Env epitopes to neutralizing antibodies by modulating Env structural conformation (35, 36, 40). We hypothesized that if P2X1 antagonists bind directly to Env, their inhibitory potencies would be modulated by the ΔCT mutation. Dotted lines indicate inhibition of NL-CI infection based with IC_50_ values based on Figure 1 and Table 1. We observed that NF279 inhibition was shifted from an IC_50_ of 1.0 μM for NL-CI to 1.8 μM for NL-CIΔCT (Figure 3B). NF449 inhibition was shifted from an IC_50_ of 0.2 μM to 2.7 μM, nearly 5-fold. By contrast, no appreciable difference in inhibition was noted for A430879. The NF279 and NF449 data demonstrate altered inhibition of HIV-1 infection that results from the conformational changes of Env induced by the ΔCT mutation, indicating that these P2X1 antagonists confer HIV-inhibitory activity by directly interacting with Env.

### NF449 inhibits a strain of CXCR4-tropic virus more efficiently than a strain of CCR5-tropic virus in primary cells

We next evaluated the impact of these drugs on inhibition of viral replication on various types of Envs to understand the spectrum of inhibition of these drugs, not only against laboratory-adapted strains but also clinical isolates based on prior observations that non-selective inhibitors could reduce both R5 and S4-tropic virus (23). To test physiological relevance, we compared inhibition of these viruses in human peripheral blood mononuclear cells (PBMCs), and in particular, we looked at transmitted/founder (T/F) viruses. Transmitted/founder (T/F) viruses arise from the propagation of a single virus strain in a naïve host and represent the variants transmitted in human HIV-1 infections We first tested the effect of P2X1 antagonists on replication of the laboratory-adapted HIV-1 NL4-3 (NL-CI, X4-tropic) virus vs. RHPA (NL-CI-11036, T/F R5-tropic The inhibition of NF449 and A438079 was compared to PPADS and AZT (Figure 4). In the presence of the X4-tropic virus, AZT inhibited productive infection to near 90%, PPADS inhibited infection to near 60%, NF449 inhibited near 75%, and A438079 inhibited 35%, which is lower than 50% inhibition observed for MT4 cells (Figure 1). For R5-tropic viral inhibition, the drugs targeting P2X receptors had decreased inhibitory activity when compared to X4-tropic viral inhibition. NF449 had an IC_50_ of 33 μM and a maximal inhibition near 45% as compared with PPADS with an IC_50_ at 21 μM and a maximal inhibition near 50%. A438079 failed to inhibit either virus above 50%, even at maximal concentrations. By contrast, AZT showed no significant change in inhibitory activity between X4- and R5-trophic infection of human PBMCs. We concluded that NF449 is an effective inhibitor of productive infection in primary cells with inhibition favoring X4-tropic viral infection.

**Figure 4.**
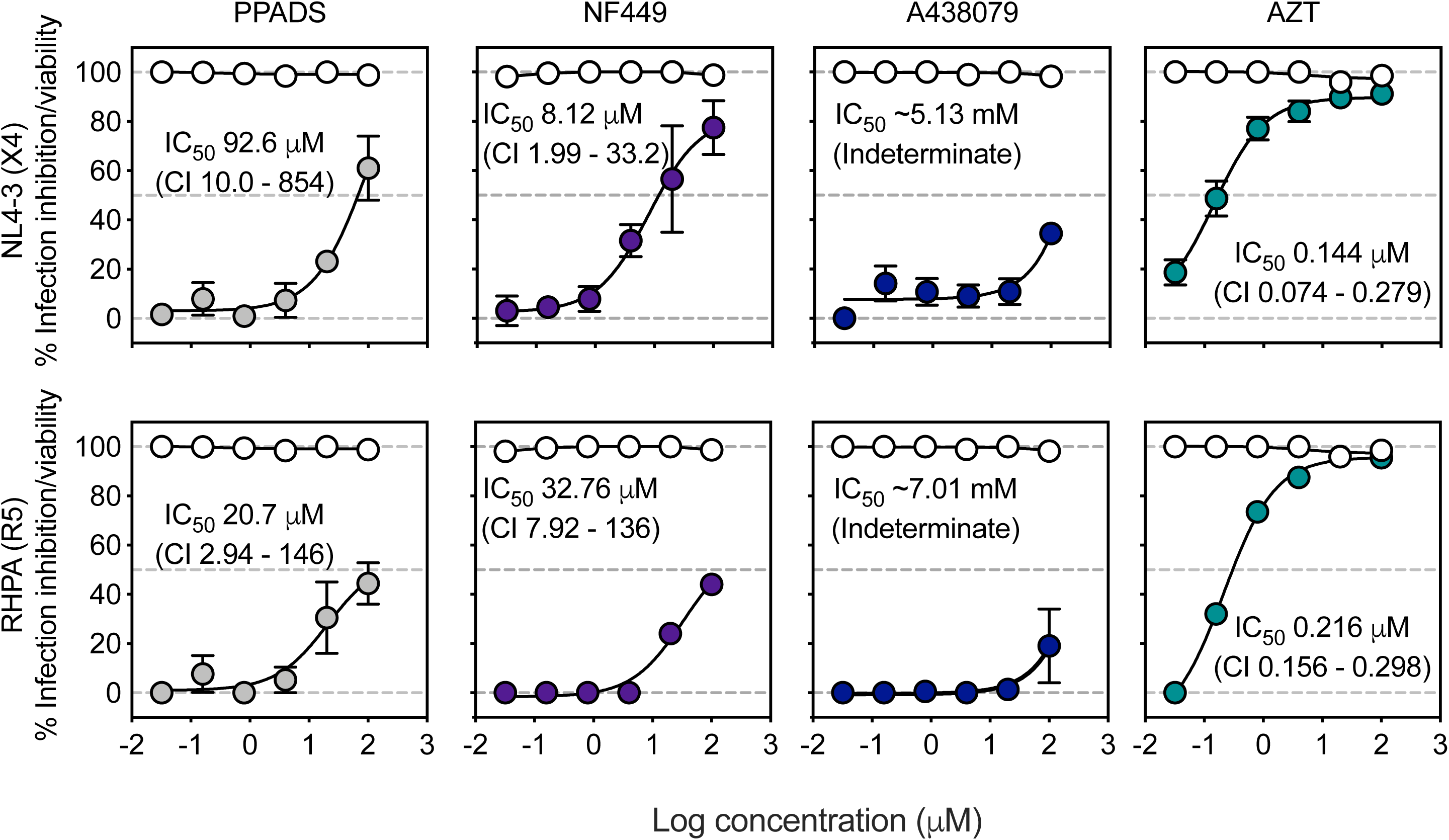
NF449 blocks HIV-1 productive infection in human PBMCs and tonsils in a strain of CXCR4-tropic and CCR5-tropic virus. PBMCs were infected with HIV-1 NL4-3 (NL-CI, X4 tropic) or RHPA (NL-CI-RHPA, R5-tropic) in the presence of indicated inhibitor. Non-linear regression curve fits are shown. Infections were conducted in the presence of 5-fold dilutions of inhibitor from 100 μM, and samples were incubated for 48 hrs and then fixed and quantified based on mCherry fluorescence. Inhibition is calculated based on the infection in the presence of drug as a function of the infection for each virus without inhibitor. Viability is plotted in the unfilled circles and percent inhibition is plotted in the black circles. Results are the means ± SEMs of at least three independent experiments.

### NF279 and NF449 inhibition is dependent on Env sequence

Next, we tested the P2X1 antagonists for inhibition of infection by HIV-1 expressing different Env variants, including T/F Env strains. We hypothesized that if the P2X1 selective antagonists do interact with Env during HIV-1 infection, as our PBMC data suggest (Figure 4), then they would have varying inhibitory activity against viruses with different Env strains. TZM-bl cells were incubated in the presence of NF279 and NF449 as compared with the less effective A438079 and tested for the ability of P2X1 antagonists to block infection of Tier 1 and Tier 2 viruses expressing various HIV-1 envelopes (Figures 5A and 5C): NL4-3 (Clade B, lab-adapted chronic X4-tropic, Tier 1), RHPA (Clade B, T/F R5-tropic, Tier 2), REJO (Clade B, T/F R5-tropic, Tier 2), SF162 (Clade B, chronic R5-tropic, Tier 1), DU172 (Clade C, T/F R5-tropic, Tier 2). These were selected to allow for a comparison of chronic and transmitted founder viruses of various clades. NF279 inhibited all viruses to 100%, with IC_50_ values between 0.04 μM and 50 μM. Of note, NF279 was remarkably potent against DU172, a molecular clone that is highly resistant to broadly neutralizing antibodies (Figures 5A and 5C). NF449 inhibited all viruses, except SF162, to nearly 100% at 100 μM, with IC_50_ values ranging from 3 μM to >30 μM across clades. A438079 had the least effective inhibition profile with a maximal inhibition of NL4-3 to 40% at 100 μM. Additionally, we looked at the effect of reducing multiplicity of infection (MOI) for the different viruses when treated with NF279 and NF449. For infection with NL4-3, RHPA, REJO, and DU172, no difference was noted in inhibition with varied MOI. For SF162, inhibition by NF279 and NF449 was greatest at low MOI whereas with increased MOI, inhibition was reduced. These data demonstrate the drug potencies vary across different viral Env constructs. This suggests that P2X1 antagonist activity is Env strain-dependent.

**Figure 5.**
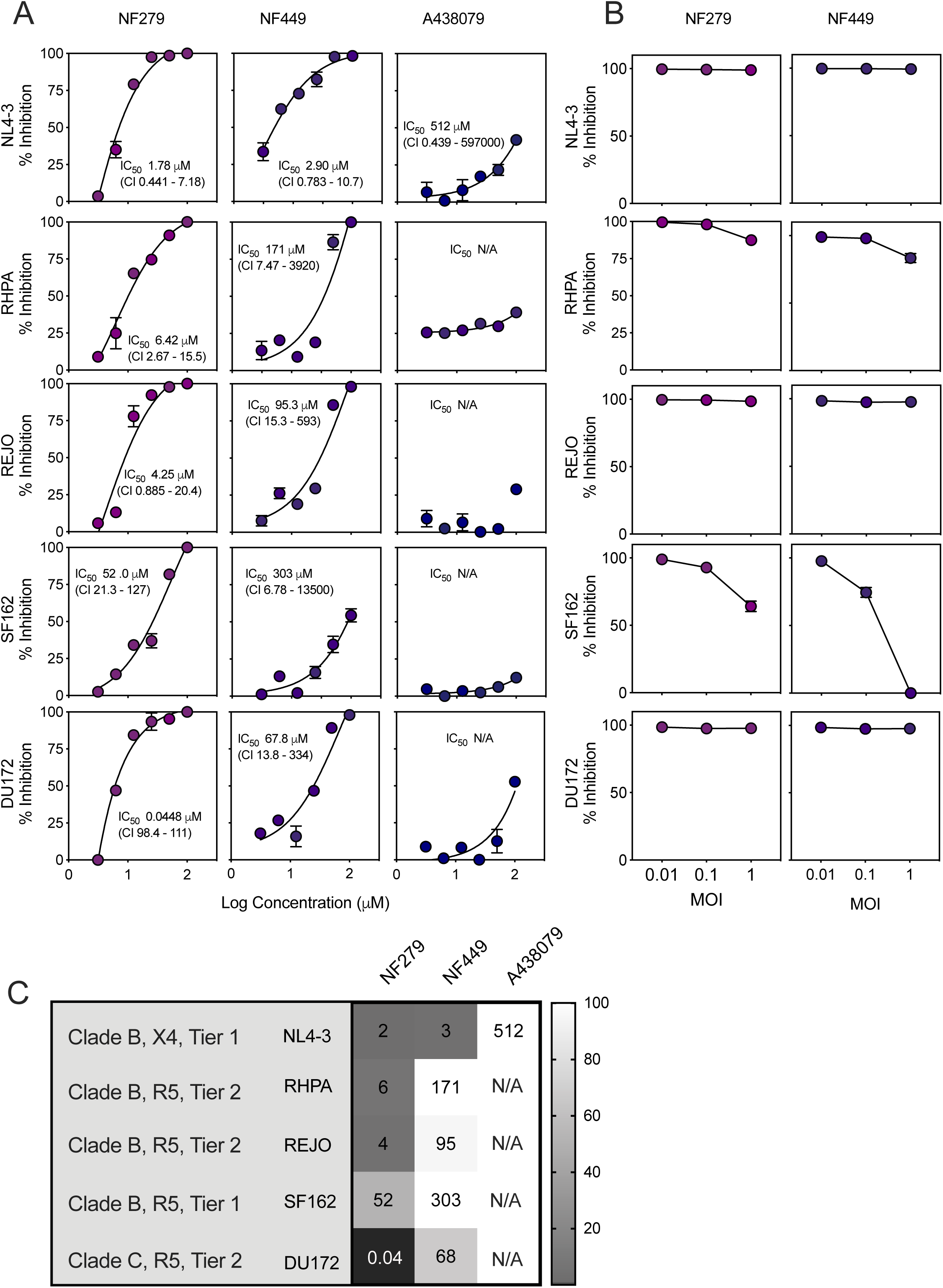
NF279 and NF449 reduce infection of multiple HIV-1 envelopes. (A) TZM-bl cells were infected with HIV-1 NL-4-3 as compared to recombinant molecular clones carrying primary *env* genes in cis: RHPA, REJO, SF162, and DU172 with dose-response curves the presence of inhibitors NF279, NF449, and A438079. Nonlinear regression curve fits for cell-free infection are shown and the calculated IC_50_ values with 95% confidence intervals indicated. Infections were conducted in the presence of serial 2-fold dilutions of inhibitor from 100 μM, and samples were incubated for 48 hrs, fixed and analyzed by flow cytometry. Inhibition is calculated based on the infection in the presence of drug as a function of the infection for each recombinant virus without inhibitor. (B) TZM-bl cells were infected with the HIV-1 recombinant Env from NL4-3, RHPA, REJO, SF162, and DU172 at multiplicity of infection (MOI) values of 0.01, 0.1, and 1 in the presence of 100 μM of either NF279 or NF449. (C) IC_50_ values were calculated from the IC_50_ values indicated in Figure 4A and plotted in a heatmap with the indicated viruses and their clade and tropism designation. Results are the means ± SEMs of at least three independent experiments.

### NF449 inhibition interferes with a V1V2-specific monoclonal antibody against HIV-1 Env

Given the observation that variations in HIV-1 Env sequence altered NF279 and NF449 inhibition, we wished to determine the region of HIV-1 envelope likely to be involved in this inhibition by looking for functional interactions with the activity of a panel of broadly neutralizing antibodies (bNAbs) (Figure 6A). We tested bNAbs targeting several regions of HIV-1 envelope (see Figure 6B), 2F5 (MPER on gp41), PG9 (gp120 V1V2 apex), 2G12 (gp120 glycan), and VRC01 (gp120 CD4-binding site). mCherry-expressing NL-CI virus was treated with drug (NF279, NF449, or AZT) followed by bNAb, and then added to MT4 cells.

**Figure 6.**
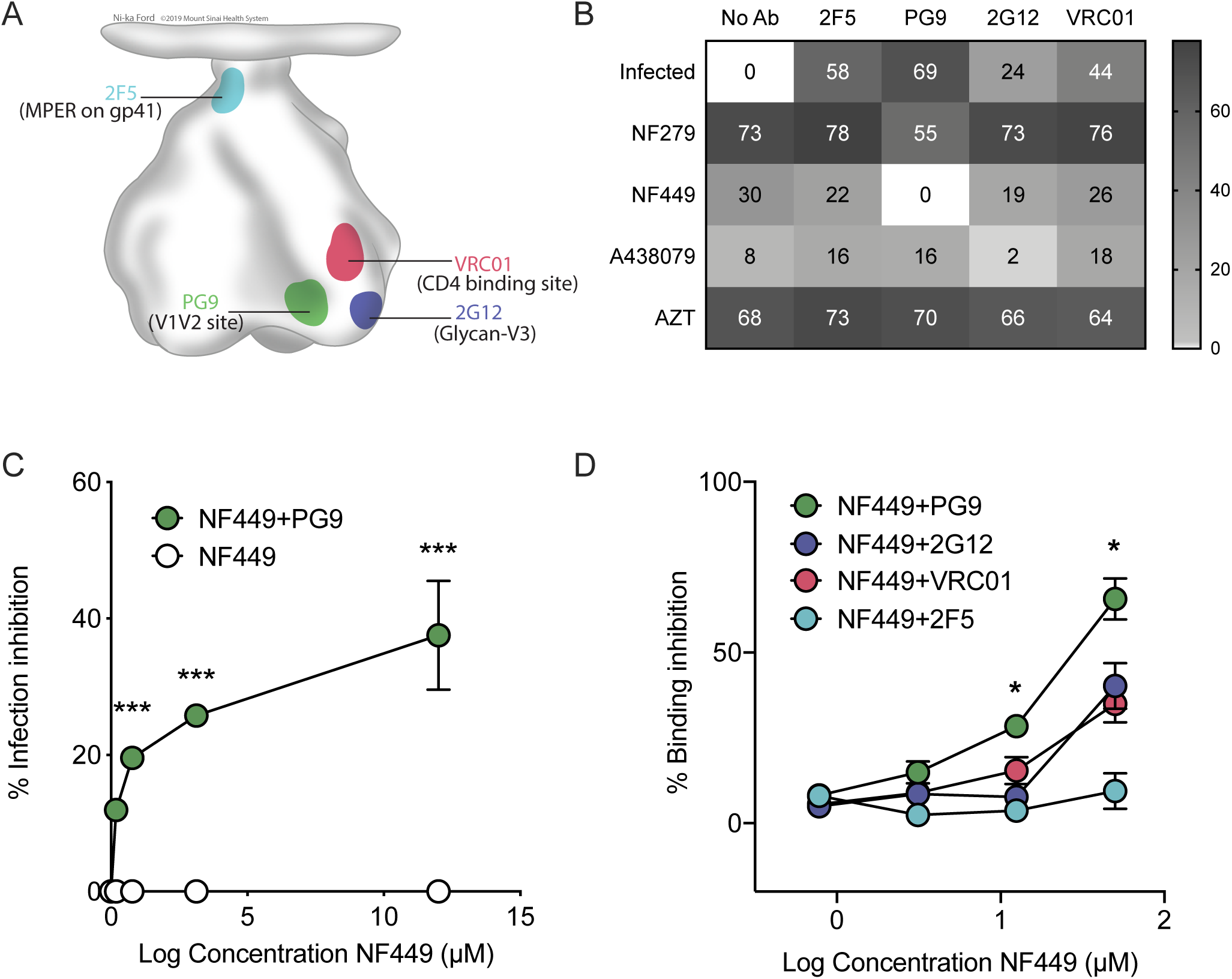
Cross inhibition by NF279 and NF449 reduces neutralization and binding by bNAb PG9. (A) A model of HIV-1 Env indicates the binding sites for each of the four bNAbs: PG9 corresponds to V1V2, 2F5 corresponds to MPER on gp41, 2G12 corresponds to Glycan-V3, and VRC01 corresponds to the CD4 binding site. (B) NF279, NF449, A438079 and AZT were tested for the ability to modify the neutralization by four bNABs: PG9, 2F5, 2G12, and VRC01. Samples were pre-treated with sub-IC_50_ inhibitory concentrations of indicated drug (0.25 µM) and/or bNAb (PG9 0.36 µg/ml, 2F5 0.06 µg/ml, 2G12 0.1 µg/ml, 0.17 µg/ml) for 30 minutes prior to addition of virus (1.62 ng of HIV-1 NL-CI). Samples were incubated for 48 hrs, fixed and analyzed by flow cytometry. Results are the means ± SEMs of three independent experiments. A heatmap of infection inhibition is shown. (C) Titrations in the presence or absence of serial 5-fold dilutions of inhibitors from 100 μM NF279 (left) and NF449 (right) are shown. Green filled circles indicate the drug with a fixed concentration of PG9 (0.36 µg/ml) and unfilled circles indicate the drug alone. Inhibition is calculated based on the infection in the presence of drug as a function of the infection without inhibitor. Nonlinear regression curve fits are shown. (D) A cell based Tetherin-high antibody binding assay is represented as % inhibition of antibody binding in the presence of increasing concentrations of NF449. Results are the means ± SEMs of at least three independent experiments.

Each of the drugs and bNAbs was tested at a concentration close to their respective IC_50_. After 48 hours, cells were fixed and analyzed by flow cytometry to quantify infection based on mCherry expression. We hypothesized that if a drug interferes with bNAb binding to Env, virus inhibition will be dampened relative to that observed with a non-interfering drug. A heatmap is shown (Figure 6B) demonstrating the inhibition of infection by drug and/or bNAb as compared to untreated infected cells. While it was expected that viral inhibition would be reduced with a non-interfering drug, it was observed that treatment with PG9 and NF449 resulted in full abrogation of inhibition. This suggested that NF449 antagonized the PG9 binding site, gp120 V1V2. Furthermore, titrations of NF449 at a fixed dose of PG9 (Figure 6C) demonstrated higher levels of infection in cells treated with both NF449 and PG9, supporting the notion that NF449 may block the ability of PG9 to bind to HIV-1 Env, either directly or indirectly through interaction and modification of the PG9 binding site, and that this interaction may account for the ability of NF449 to inhibit viral membrane fusion.

To further explore the hypothesis that NF449 activity is dependent on directly binding to, or changing the conformation of the HIV-1 Env V1V2 region at the PG9 binding site, we employed a well-characterized antibody binding system (41-43). This system retains viral particles on the surface of HIV-infected CD4 T cells based on tetherin-high expression, allowing for the enhanced resolution of differences in antibody binding. We incubated HIV-1 infected cells at 37°C in the presence of increasing concentrations of NF449. Prior to conducting binding experiments, residual NF449 was washed off and Ab binding experiments were conducted at 4°C. We observed that PG9 binding to HIV-1 Env was significantly reduced in the presence of increasing concentrations of NF449, as compared to binding of 2G12, VRC01, and 2F5 binding (Figure 6D). The highest level of inhibition was observed at 50 μM where PG9 binding inhibition was noted at 66% as compared to 2G12 inhibition of 41%, VRC01 inhibition of 33%, and 2F5 inhibition of 9%.

Taken together, these observations demonstrate modulation of NF449 fusion inhibition by Env-specific bNAbs and differing Env variants, suggesting that Env plays a role in P2X1 antagonist inhibition of HIV-1 infection. The observation that NF449 and PG9 antagonize inhibition suggests that NF449 may binding directly to gp120 at the region of V1V2 or allosteric changes to the Env structure may prevent PG9 access to Env V1V2. These may impact on interaction with coreceptor critical to mediate viral membrane fusion.

## Discussion

Here we demonstrate that several P2X1 antagonists can block HIV-1 infection, and that the inhibition (1) interferes at the level of viral membrane fusion (2) favors X4-tropic virus (3) relies on intact gp41 and (4) demonstrates varied activity against diverse clades. This activity may specifically interfere with Env, as NF279 and NF449 interfere with antibody accessibility at the V1V2 region of gp120. In Figure 1, it was demonstrated that NF279 and NF449 inhibit infection at micromolar concentrations, with IC_50_ values lower than for the non-selective inhibitor PPADS. A438079 had minimal inhibition and the other drugs tested did not interfere with HIV-1 productive infection.

NF279 and NF449 inhibited viral membrane fusion, consistent with our prior observations (25). A time-of addition experiment indicated that both NF279 and NF449 both inhibit HIV-1 replication before 4 HPI, concurrent with AMD3100 inhibition but distinct from AZT activity that works later in the virus life cycle. These data further support the notion that these P2X1 antagonists block HIV-1 infection at the level of viral membrane fusion. It is important to note that NF279, NF449 and A438079 may have activity against both P2X1 and P2X7. NF279 and NF449 have greater selectivity for P2X1 and A438079 has greater selectivity for P2X7 (44-51); however, the concentrations tested in this study are much higher than the range where they act as selective antagonists, and therefore, the observations cannot definitively distinguish the affected receptor subtypes (Table 1). It is important to note that even at the highest concentrations, viability of cells was not affected.

A cross titration experiment was performed to determine whether treatment with enfurvutide (T-20) inhibition would be impacted by NF279 or NF449 treatment as the mechanism of T-20 activity is based on binding of the pre-hairpin structure of gp41 to inhibit fusion (Figure 3A). We reasoned that if the drugs have similar mechanisms, coordinated treatment with T-20 and NF279 or NF449 would competitively antagonized inhibition of HIV-1 infection. Our observations did not support this as no difference was noted between NF279 or NF449 treated samples and those treated with NF279 or NF449 plus T-20. These data suggest that NF279 and NF449 may have an alternative mechanism to engage Env. We then tested the hypothesis that HIV-1 Env conformation might play a role in virus fusion inhibition by these drugs by evaluating the effect of a C-terminal truncation mutant of gp41 (NL-CIΔCT) that open the HIV-1 Env structure to expose neutralizing Ab epitopes. Virus inhibition was shifted slightly for NF279 but nearly 5-fold for NF449, indicating higher resistance of altered Env exposure to inhibition by NF449 (Figure 3B). This observation may be explained by a different conformation of Env that is induced by the gp41 mutation to result in altered binding affinity of the drugs, which is demonstrated by a change in the slope of NF449 inhibition curves against NL-CI versus NL-CIΔCT (52). A438079 demonstrated no change in its inhibition against NL-CI compared to NL-CIΔCT, suggesting that the effects seen with NF279 and NF449 are specific to these two drugs (52).

To assess HIV-1 inhibition in physiologically relevant cells, we compared NF279, NF449, and A438079 were to AZT for the ability to inhibit HIV-1 productive infection in human PBMCs (Figure 4), We observed inhibition by both NF279 and NF449, with greater activity of NF449 against of X4-tropic virus than R5-tropic virus, and minimal overall activity of A4388079. NF279 and NF449 were also compared to A438079 across a variety of HIV-1 envelopes (Figure 5). NF279 maintained the lowest IC_50_ values of the three compounds with only modest increases in IC_50_ between NL4-3 and RPHA envelope. The CXCR4-tropic virus was more sensitive to NF449 inhibition than CCR5-tropic virus, which suggests envelope specificity; however, NF279 and NF449 inhibited a range of envelopes suggesting that the nature of the inhibition may be through direct binding to Env. It is notable that NF449 inhibited SF162 less than other viruses, while NF279 inhibited DU172 to a much greater extent. Neutralization of SF162 by NF449 may relate to Env variable loop 2 (V2) glycosylation and interference with CD4 and co-receptor interactions (53). Highly effective inhibition of DU172 by NF279 suggests that these compounds may be potential therapeutic options for highly resistant viruses. In Figure 5B, MOI titrations indicate minimal change in inhibition in all viral clones with the exception of NF449 in which increasing the MOI to 1 reduces infection nearly completely with NF449, suggesting the ability of that clone to compete out the effect of inhibition.

Based on evidence that varied HIV-1 Env clones had different responses to NF279 and NF449 inhibition, we sought to map the region of inhibition by testing the effect of treatment with NF279 and NF449 on of a panel of bNABs for their ability to inhibit HIV-1 infection (Figure 6). Our observation indicated that PG9 inhibition was significantly reduced in the presence of NF449 in a dose-dependent manner. These observations suggest that NF449 may inhibit HIV-1 Env at the region of V1V2 where PG9 binds. Furthermore, the co-administration of NF449 and PG9 negated the effect of inhibition, suggesting an allosteric interaction at the site of V1V2 or director interaction between drug and antibody, reducing effective binding and neutralization of HIV-1.

We observed in an *in vitro* system, NF279 and NF449 but not A438079, can effectively block HIV-1 replication at the level of viral membrane fusion (Figure 2). Both compounds are most effective at inhibiting X4-tropic HIV-1 infection, but are effective inhibitors against multiple HIV-1 envelopes. However, NF279 and NF449 are suramin-derivatives, which are large molecules that may not be optimal for therapeutic development for the following reasons. Therapeutic options that target P2X receptors for pharmaceutical development would include those agents that follow Lipinski’s parameter thresholds (54, 55): molecular weight of less than 500 kDA, lipophilicity (octanol-water partial coefficient) logP <5, no more than 10 hydrogen bond acceptors, and no more than 5 hydrogen bond donors. The values for IC_50_ reported in Figure 1 are higher than those IC_50_ values reported for P2X1 inhibition (44-46, 49-51, 56-69). This may reflect that the drug concentrations tested have effects that target not only the receptor but other cellular events such as the direct binding of HIV-1 Env. It will be of benefit to test more compounds with P2X1 and P2X7 activity that satisfy these pharmaceutical requirements for potential clinical development. By contrast to NF279 and NF449, A438079 did not demonstrate inhibition of HIV-1 fusion. Interestingly, A438079 has previously been demonstrated to have differential inhibition of HIV-1 productive infection in cell lines vs. in human lymphoid aggregate cultures (24). This may reflect a different mechanism of inhibition from the P2X1 antagonists and will be the subject of future investigation.

Giroud et al. (22) have demonstrated that NF279 can antagonize signaling of CCR5 and CXCR4 and conclude that this drug and other drugs function to inhibit HIV-1 fusion through interferences with functional engagement of CCR5 and CXCR4 by HIV-1 Env. These studies indicate inhibition of fusion through co-receptor binding and through antagonism of calcium signaling stimulated by gp120. Several studies implicate the role of V1V2 in CXCR4 and CCR5 interaction (37, 70-72). In an uninhibited setting, HIV-1 fusion occurs through Env, a trimeric glycoprotein with three copies of the gp120 subunit and three copies of the membrane-anchored gp41 subunit. The gp120 subunit binds to the host cell CD4 which induces conformational changes that expose the binding site of coreceptor (CCR5/CXCR4) and this mobilizes the gp41 fusion peptide which anchors in the host cell membrane to establish fusion. A model is shown in Figure 7 in which HIV-1 Env is exposed and can bind to CD4, recruit coreceptor, and mediate viral membrane fusion. We propose that treatment with NF449 results in binding of HIV-1 Env V1V2 which interferes with exposure of the binding site for coreceptor engagement. The model demonstrates an effect in which NF449 may binding directly to V1V2 which may result in failure to recruit receptor (Figure 7B1). Alternatively, NF449 may bind to another site, resulting in conformational change that limits V1V2 access by PG9 (Figure 7B2). When NF449 or NF279 come in contact with HIV-1 Env at the V1V2 loop, a conformational change or steric hindrance prevents the binding of HV-1 Env to recruit coreceptor, leading to a failure to complete HIV-1 fusion. When NF449 is added together with PG9, there is abrogation of the inhibition caused by PG9 or NF449 alone. The inhibition profiles for NF449 vary and will help to elucidate the development of targeted drugs that can serve to inhibit HIV-1 Env and block HIV-1 viral membrane fusion. Future studies will address whether escape mutants of NF449-treated cells develop Env mutations in the V1V2 site and if mutations in the V1V2 region are resistant to NF449 inhibition.

**Figure 7.**
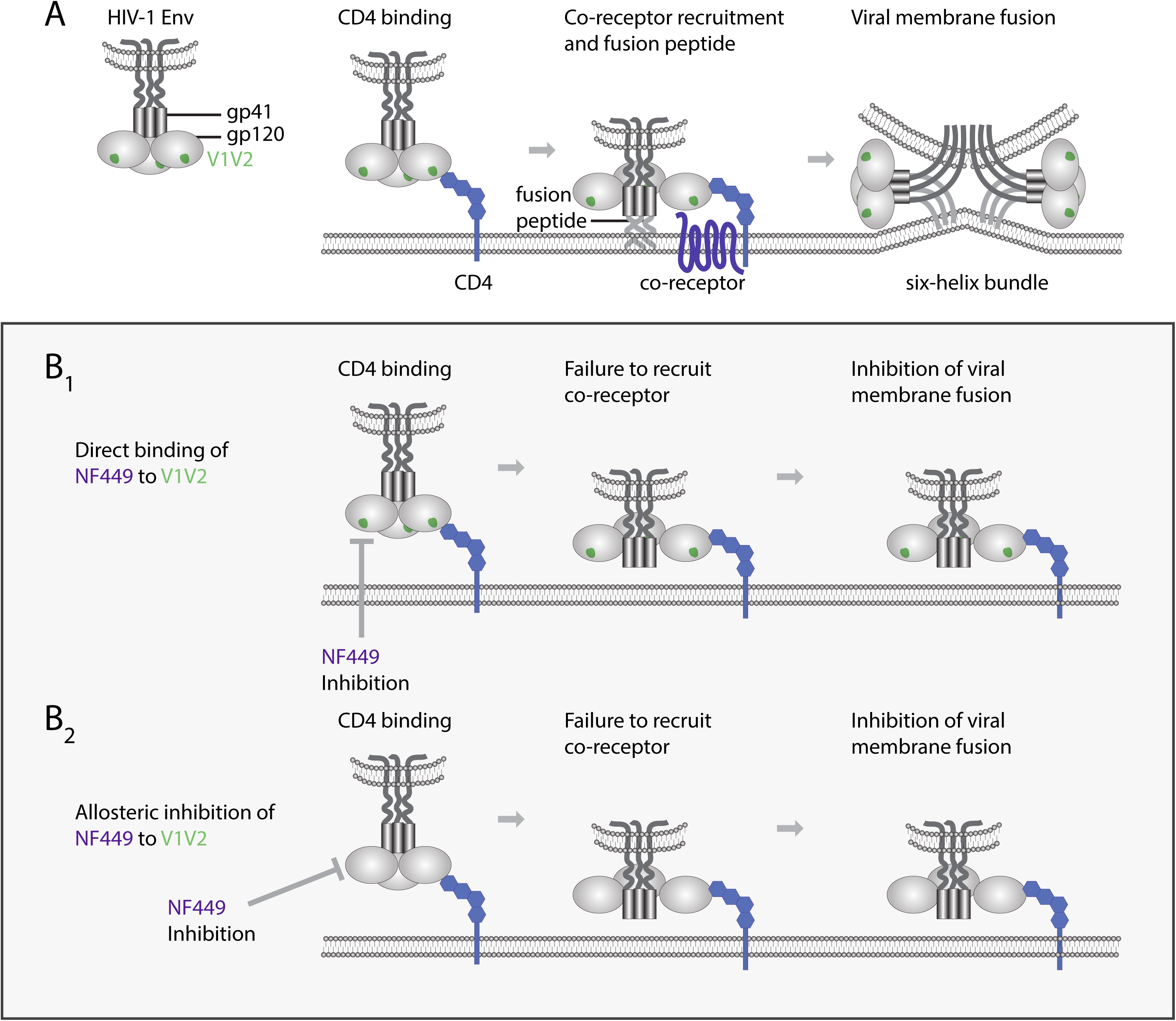
Model of NF279 and NF449 inhibition of HIV-1 Env. HIV-1 is shown with the gp120 V1V2 region in green. (A) Binding of gp120 to CD4 enables coreceptor recruitment and fusion peptide insertion which leads to six-helix bundle formation and viral membrane fusion. (B1) NF449 binding directly to gp120 V1V2 results in inability to recruit CCR5/CXCR4 and facilitate viral membrane fusion. (B2) NF449 binding to an alternative site results in conformational change which reduces access of V1V2 and results in inability to recruit CCR5/CXCR4 and facilitate viral membrane fusion.

Based on this model, we propose that P2X1 antagonists may represent a novel adjunctive therapeutic strategy whereby drug binding to HIV-1 Env can cause a conformational change or steric hindrance, preventing coreceptor recruitment and viral membrane fusion that is negated in the presence of the anti-V1V2 antibody PG9. We conclude from these studies that P2X1 antagonists may represent potential HIV-1 therapeutic options that serve to inhibit cross-clade HIV-1 replication. Further studies will be necessary to identify selective inhibitors that are amenable to pharmacologic development and define the precise mechanism of inhibition. Regardless, these observations introduce important prospects for dually active therapeutic options that would reduce the burden of morbidity and mortality of chronic inflammation in HIV-1-infected individuals.

## Materials and Methods

### Cells and cell lines

The human cell line MT4 (provided by Douglass Richman, NIH AIDS Reagent Program (ARP)) cells were maintained in RPMI 1640 medium (Sigma) which contained 10% fetal bovine serum (FBS; Sigma), 100 U/ml penicillin (Gibco), 10 U/ml streptomycin (Gibco), and 2 mM glutamine (Gibco) (complete RPMI). Pseudoviruses were produced with 293T/17 cells (American Type Culture Collection, ATCC). Neutralization assays used the TZM-bl cell line used as target cells, obtained from Dr. John C. Kappes, Dr. Xiaoyun Wu, and Tranzyme Inc. through the NIH ARP. Both 293T/17 and TZM-bl cells were maintained in Dulbecco’s modified Eagle Medium (DMEM; Sigma) containing 10% cosmic calf serum (CCS; HyClone) and 100 U/mL penicillin, and 10 U/mL streptomycin 2 mM glutamine (Gibco) (complete DMEM). PBMCs were obtained from de-identified HIV-1 negative blood donors (New York Blood Center), purified by Ficoll (HyClone) density gradient centrifugation and were maintained in RPMI 1640 medium (Sigma) containing 10% fetal bovine serum (FBS; Sigma), 100 U/ml penicillin (Gibco), 10 U/ml streptomycin (Gibco), and 2 mM glutamine (Gibco) (complete RPMI).

### Plasmids and viruses

Plasmids expressing HIV-1 Env of 6535 (clone 3 (SVPB5)), BaL (clone BaL.01), REJO (pREJO4541 clone 67 (SVPB16)), and ZM109 (ZM109F.PB4, SVPC13) were used for generating HIV-1 pseudoviruses. These plasmids were obtained from Drs. David Montefiori, Feng Gao, Ming Li, John Mascola, B.H. Hahn, J.F. Salazar-Gonzalez, C.A. Derdeyn and E. Hunter through the NIH ARP. The plasmid bearing SF162 (pIRESSF162) *env* was constructed as previously described (73, 74). HIV-1 NL-GI contains green fluorescent protein (GFP) in place of *nef*, and *nef* expression is directed by a downstream internal ribosome entry site (IRES) (28). HIV-1 NL-CI contains mCherry in place of *nef*, as above. PBMCs were infected with NL-GI/NL-CI, which contains the NL4-3 envelope (X4 tropic), or RHPA, which was constructed by insertion of the R5-tropic B-clade primary envelope from pRHPA4259 clone 7 (SVPB14) into NL-GI (75). The gene for the RHPA clone was obtained, from B. H. Hahn and J. F. Salazar-Gonzalez (ARP). HIV-1 Gag-iCre is an HIV-1 clone carrying Cre recombinase as a Gag-internal gene fusion that releases active Cre into cells upon viral entry activating a recombinatorial gene switch changing dsRed to GFP-expression (25). NL-CIΔCT was cloned by generating a PCR fragment of the C-terminal Env from the NL-CIΔCT plasmid as described (36). Pseudoviruses were produced by co-transfecting 293T/17 cells with HIV-1 *rev-* and *env-*expressing plasmids and the pNL4-3Δenv R-E-plasmid using the jetPEI transfection reagent (Polyplus-transfect SA). Supernatants were harvested after 48 hours and clarified by high-speed centrifugation (Sorvall ST 40R Centrifuge, Thermo Fisher Scientific) at 100,000 x g at 4°C for 2 hours and 0.45 μm filtration. Viral stocks were quantified by the HIV-1 p24 antigen via enzyme-linked immunosorbent assay (ELISA) with coating antibody D7320, sheep anti-HIV-1-p24 gag (Aalto Bio Reagents) as described previously (36). Single-use aliquots were stored at −80°C.

### Antagonists

A panel of antagonists was tested using 5-fold serial dilutions beginning at 100 μM, unless otherwise stated. These included PPADS (Sigma), A438079 (Tocris), NF449 (Tocris), NF279 (Tocris), JNJ479655567 (Tocris), A804598 (Tocris), A839977 (Tocris), A740003 (Tocris), AZ10606120 (Tocris), AZ11645373 (Tocris), GW79134 (Tocris), JNJ4795567 (Tocris), and AZT (Sigma), T-20 (0.1 μg/ml, Sigma).

### Broadly neutralizing monoclonal antibody (bNAbs)

A panel of bNAbs targeting different epitopes on HIV-1 Env were tested for their ability to block HIV-1 infection and binding. These include: anti-V1V2 Apex MAb PG9 (International AIDS Vaccine Initiative [IAVI], ARP) (200 μg/ml); anti-gp41 MAb 2F5 (International AIDS Vaccine Initiative [IAVI], ARP) (1 mg/ml); anti-gp120 glycan MAb 2G12 (Hermann Katinger, ARP) (100 μg/ml); and VRC01 (International AIDS Vaccine Initiative [IAVI], ARP) (3 mg/ml).

### Flow cytometry and gating strategy

An LSR II flow cytometer (BD Biosciences) was used to detect infection and discriminate donor and target cell populations. Viable cells in productive infection assays were detected with LIVE/DEAD Fixable Dead Cell Stain (Life Technologies), an amine reactive fluorescent dye that can penetrate the membranes of dead cells but not live cells. Samples were stained with LIVE/DEAD Fixable Blue Dead Cell Stain or LIVE/DEAD Fixable Violet Dead Cell Stain at a concentration of 1:1000 in Wash Buffer (PBS supplemented with 2 mM EDTA and 0.5% bovine serum albumin). Stained cells incubated at 4°C for 30 minutes, then were washed and fixed in 2% paraformaldehyde for flow cytometry. All cells were initially discriminated by side scatter (SSC) area versus forward scatter (FSC) area (SSC-A/FSC-A); doublets were excluded using FSC height (FSC-H) vs FSC-A and dead cells were excluded by gating on the negative populations for LIVE/DEAD Fixable Dead Cell Stain. In productive infection assays, infection was detected by the presence of mCherry in cells infected with HIV-1 NL-CI or the presence of GFP in cells infected with HIV-1 NL-GI. GFP was detected using the fluorescein isothiocyanate (FITC) channel, dsRed-Express and mCherry were detected using the phycoerythrin-Texas Red (PE-Texas Red) channel, LIVE/DEAD Fixable Violet Dead Cell Stain was detected with the 3-carboxy-6,8-difluoro-7-hydroxycoumarin (Pacific Blue) channel, and LIVE/DEAD Fixable Blue Dead Cell Stain was detected with the 4’,6-diamidino-2-phenylindole (DAPI) channel. In HIV-1 fusion assays, donor cells labeled with eFluor 450 were detected in Alexa Fluor 405-A channel and target cells that express dsRed were detected using Alexa Fluor 568-A channel. Transfer of gag-iCre from donor to target cells was measured on the basis of activity of Cre Recombinase, which results in recombination of Lox P sites switching the phenotype of the targets to GFP positive. This switch from dsRed to GFP was measured using Alexa Fluor 568-A channel versus Alexa Fluor 488-A channel. All cells within a single experiment were analyzed using the same instrument settings.

### Productive infection assays

Target MT4 cells were infected in 96-well plates with HIV-1 NL-GI/NL-CI/NL-CIΔCT/NL-CI-RHPA (1.62 ng p24 per well) to obtain up to 10% infection after 48 hours in the absence of inhibitors. MT4 cells were pre-incubated with antagonists for 30 min at 37°C before infection with HIV-1. At 48 hours after infection, cells were fixed in 2% paraformaldehyde and infection was quantified via GFP or mCherry fluorescence in flow cytometry. For the time-of-addition assay, P2X1 antagonists were added to MT4 cells at the indicated time points of HPI with HIV-1 NL-CI (1.62 ng p24 per well), as previously described (29). At 48 hours after mixing, cells were stained and fixed in 2% paraformaldehyde for flow cytometry, as described above. For coreceptor competition assays, MT4 cells were co-incubated with P2X antagonists and HIV-1 fusion antagonists for 30 min at 37°C before infection with HIV-1 NL-CI (1.62 ng p24 per well). At 48 hours post infection, cells were stained and fixed in 2% paraformaldehyde for analysis via flow cytometry as described above.

### Virus Tropism in PBMCs

PBMCs were activated with PHA (4 μg/ml) and IL-2 (50 U/mL) for 3 days and infected by spinoculation, as previously described (29). Briefly, 2.5 × 10^5^ cells were incubated in the presence or absence of indicated inhibitors in a 96-well flat bottom plate for 30 minutes at 37°C then spun at 1,200 x *g* for 99 minutes with 47.7 ng HIV-1 NL-CI or NL-CI RHPA. After overnight incubation at 37°C, the culture medium was replaced with complete RPMI containing IL-2 (50 U/ml) and 10 μM AZT. At 48 h after spinoculation, cells were stained and fixed in 2% paraformaldehyde for flow cytometry, as described above.

### HIV-1 fusion assay

A stable cell line of Jurkat cells called Jurkat floxRG (where RG indicates red to green) was used as target cells and was generated by retroviral transduction with the pMSCV-loxP-dsRed-loxP-eGFP-Puro-WPRE vector (Addgene, plasmid 32702) (76), which expresses a dsRed reporter flanked by *loxP* sites followed by a Cre-activated enhanced GFP (eGFP) gene (76). Jurkat (donor) cells were transfected by nucleofection (Amaxa Biosystems) with 5 μg HIV-1 Gag-iCre DNA, cultured overnight in antibiotic-free medium, and purified by Ficoll-Hypaque density gradient centrifugation. Target cells were detected on the PE-Texas Red channel, and donor cells were preincubated separately with P2X1 antagonists for 30 min at 37°C before mixing 1.25 × 10^5^ cells at a ratio of approximately 1:1, co-cultured at 37°C for 48 hours, and fixed, as described previously (25).

### Infectivity and Neutralization assay

Virus infectivity and neutralization was measured using a β-galactosidase-based luciferase assay (Promega) with TZM-bl target cells, as previously described (75). Briefly, serial dilutions of antagonists were added to the virus in 96-well plates (Costar) and incubated for the designated time period at 37°C. TZM-bl cells were then added with or without DEAE-dextran (6.25μg/ml; Sigma). After incubation for 48 hours, a luciferin-galactoside substrate (6-O-β-galactopyranosyl-luciferin) was added. The cleavage of the substrate by β-galactosidase generates luminescent signals measured in RLUs. Test and control conditions were tested in duplicate or triplicate. Assay controls included replicate wells of TZM-bl cells alone (cell control) and TZM-bl cells with virus alone (virus control). Percent neutralization was determined on the basis of virus control under the specific assay condition (e.g. 1 hour or 24 hours pre-incubation of virus without mAbs). The virus inputs were the diluted virus stocks yielding equivalent RLUs (typically ∼100,000 RLUs) under the different assay conditions.

### Tetherin-high antibody binding assay

A cell-based assay for bNAb binding was used as previously described (42) in which a subclone of the Jurkat E6 CD4 T cell line that constitutively expresses high levels of tetherin, is transfected with an HIV-1 Δvpu mCherry fluorescent reporter virus. Cells retain virus particles on their surface. Cells are pre-incubated for 20 minutes with NF449 and then incubated with bNAb for 30 minutes at 4 degrees. To quantify Ab binding, a binding index (BI) was established that provides a combined measure of the frequency of opsonized HIV-infected cells (%) and the density of HIV-specific Ab binding (median fluorescent intensity [MFI]) and the results were plotted as inhibition of binding.

### Statistical analysis and calculations

Comparisons were performed using GraphPad Prism 7, version 7.0d (GraphPad Software). DMSO-treated controls were set to 100% and drug-treated conditions were expressed as a percentage of control. Statistical analyses were performed on inhibition data that reached ≥50% with a one-tailed student’s t-test. A *p* value of less than 0.05 was considered statistically significant.

## Author contributions

AYS, NDD, FP, NS, HSM, TLF and CU performed experiments. AYS, NDD, HSM, and TLF carried out productive infection assays in MT4s and PBMCs. FP and NS carried out HIV-1 Gag-iCre fusion assays. AYS performed the time-of-addition assay and coreceptor/CD4 assay. CU, CH and HSM performed and analyzed assays of TZM-bl infection of various HIV-1 envelopes. RA performed binding assays. AYS, HSM, TLF, and THS analyzed results and wrote the paper. AYS, NDD, HSM, RA, BKC, and THS designed the experiments. THS and BKC conceived the approach.

## Acknowledgements

We would like to thank the members of the Chen laboratory for meaningful discussions. T. Swartz was funded by the NIH K08AI20806 and by the Schneider-Lesser Foundation. This work was supported by grants to B. Chen from NIH, NIAID R01AI074420.

